# Sustained plumage divergence despite weak genomic differentiation and broad sympatry in sister species of Australian woodswallows (*Artamus* spp.)

**DOI:** 10.1101/2022.04.14.488308

**Authors:** Joshua V. Peñalba, Jeffrey L. Peters, Leo Joseph

**Affiliations:** Museum für Naturkunde Berlin, Leibniz Institute for Evolution and Biodiversity Science, Center for Integrative Biodiversity Discovery, Invalidenstr. 43, D-10115, Berlin, Germany; Department of Biological Sciences, Wright State University, Dayton, OH, USA; Australian National Wildlife Collection, CSIRO National Research Collections Australia, Canberra, Australia

**Keywords:** plumage divergence, sympatry, population genomics, hybridization, speciation, woodswallows

## Abstract

Plumage divergence can function as a strong premating barrier when species come into secondary contact. When it fails to do so, the results are often genome homogenization and phenotypic hybrids at the zone of contact. This is not the case in the largely sympatric masked woodswallow and white-browed woodswallow species (Passeriformes: Artamidae: *Artamus* spp) complex in Australia where phenotypic integrity is sustained despite no discernible mitochondrial structure in earlier work. This lack of structure may suggest recent divergence, ongoing gene flow or both, and phenotypic hybrids are reported albeit rarely. Here, we further assessed the population structure and differentiation across the species’ nuclear genomes using ddRAD-seq. As found in the mitochondrial genome, no structure or divergence within or between the two species was detected in the nuclear genome. This coarse sampling of the genome nonetheless revealed peaks of differentiation around the genes *SOX5* and *Axin1*. Both are involved in the *Wnt/*/μ-catenin signaling pathway, which regulates feather development. Reconstruction of demographic history and estimation of parameters supports a scenario of secondary contact. Our study informs how divergent plumage morphs may arise and be sustained despite whole-genome homogenization and reveals new candidate genes potentially involved in plumage divergence.

## Introduction

Plumage divergence between sister lineages of birds often distinguishes different subspecies and species. Often, the more divergent populations are in plumage, especially in changes of pattern rather than of simple color replacement, the more likely they are to be classified as different taxonomic species (Patten & Unitt, 2002; Paxton, 2009). This is not only because it is easier for us to distinguish between forms but also because plumage is a secondary sexual trait playing a role in mate choice and premating isolation (Mason & Bowie, 2020; Price, 2008). Large plumage differences and corresponding mate preferences can accumulate in allopatry and, when populations come into secondary contact, can serve as a strong barrier to gene flow resulting in speciation (Cowles & Uy, 2019; Sætre et al., 1997; Turbek et al., 2021). Plumage traits can also serve as a target for divergent ecological selection. If this divergence also results in differences in mate choice, plumage traits can serve as ‘magic traits’ in facilitating speciation (Servedio et al., 2011). Comparative studies on both plumage and genomic divergence have shown a positive association with plumage divergence and reproductive isolation (Winger & Bates, 2015).

Despite the expected association between plumage and genome divergence, there are also many documented cases of a lack of genomic differentiation in spite of strong plumage divergence. These unusual systems have been crucial in revealing the genetic architecture of complex plumage traits and correlative, if not causative, links between plumage divergence and particular genes involved. When most of the genome is undifferentiated or has been homogenized by gene flow, loci that remain strongly differentiated usually underlie various components of plumage traits. For example, the carrion and hooded crows (*Corvus corone*, *C. cornix*, respectively) in Europe display strong plumage integrity despite extensive gene flow in their narrow hybrid zone. An epistatic interaction between a gene, *NDP*, and a large region in chromosome 18 explains the divergence in plumage and variation in hybrid morphs (Knief et al., 2019). This genetic architecture coupled with mate-choice dynamics can explain the narrow hybrid zone despite extensive introgression (Metzler et al., 2020). Hybridization between the golden-winged warbler (*Vermivora chrysoptera*) and blue-winged warbler (*V. cyanoptera*; F_ST_ ∼ 0.0004) in North America has revealed just six divergent peaks in the genome, four of which were adjacent to genes known to be involved in feather development and pigmentation (Toews et al., 2016). The *agouti signaling protein* (*ASIP*) has been isolated as responsible for the divergence in throat coloration in the two morphs. Asymmetric introgression of the head plumage trait in two subspecies of the white wagtail (*Motacilla alba*) has revealed a partial dominant and epistatic interaction between two small regions in chromosome 1A and 20 (Semenov et al., 2021). Extensive introgression between two subspecies of the northern flicker (*Colaptes auratus*) in North America (F_ST_ ∼ 0.008) has revealed the genomic basis underlying four plumage traits and the role of *CYPJ2J19* to cause a red to yellow transition (Aguillon et al., 2021). A recent radiation in capuchino seedeaters (*Sporophila* spp) in South America resulted in nine phenotypically-distinct, largely-sympatric species (Campagna et al., 2012). Whole-genome investigation of five of the species revealed extremely low differentiation (F_ST_ ∼ 0.008) and nine melanogenesis genes in regions of high divergence, with the most fixed differences concentrated around *ASIP* (Campagna et al., 2017). Further experiments show that plumage and song differences lead to premating isolation despite the lack of population structure in the genome (Benites et al., 2015; Turbek et al., 2021).

Three common findings emerge from these studies: (1) extensive introgression can occur despite plumage divergence or, conversely, plumage divergence can be sustained despite extensive introgression; (2) one or more of many different genes can be involved in plumage divergence and so are not always shared among systems, and (3) the genes involved in plumage divergence (a secondary sexual trait) are usually found on autosomes rather than the sex chromosomes.

Another shared feature among most of these study systems except the capuchino seedeaters is that they include hybrid zones between two populations. This means that phenotypic hybrids are likely geographically restricted and can serve as a buffer for contact between the parental plumage forms. It is also likely that populations diverged in allopatry and came into secondary contact in the hybrid zone or diverged with relatively restricted gene flow only occurring in the zone of contact.

Here, we present a genomic interrogation of an unusual case that confounds these emergent themes. It concerns two Australian birds never classified as anything other than two species (see Schodde & Mason, 1999), the masked woodswallow (*Artamus personatus*) and white-browed woodswallow (*Artamus superciliosus*) (Passeriformes: Artamidae; Figure 1). Field observations and collections of museum specimens since the 1830s have established that they are largely sympatric across much of their range in the eastern half of the Australian continent just as capuchino seedeaters are extensively sympatric in South America. The same wealth of data have established that the masked woodswallow is by far the more prevalent species of the two across much of the continent’s western half except in its extreme north (Carnaby, 1965; Higgins, 2006; Johnstone & Storr, 2004; Figure S1). The two are phenotypically very different but some plumage traits are surely homologous (e.g., dark facial “mask” in both; white post-supraocular spot of masked being a vestige of the complete white superciliar stripe of white-browed; Figure 1). Especially notable given that they are a pair of closely related oscine songbird taxa is that their vocalizations appear all but identical, no discernible difference ever having been rigorously described (Higgins *et al*. 2006; *cf* Lowe and Lowe 1972). Unlike most systems cited above, however, the two are often found in mixed flocks, especially in eastern Australia, of up to hundreds of birds and breed near one another, even in the same tree (Carnaby, 1965; Higgins, 2006; Lenz, 2019). Despite this, phenotypic hybrids are rare but not unknown (Barnard, 1944; McGill, 1944); we are aware of photographic records of at least seven clearly intermediate individuals of presumed hybrid descent (e.g., see images at https://ebird.org/checklist/S79795345; https://ebird.org/australia/checklist/S61566354; Figures S2-S9).

**Figure 1.**
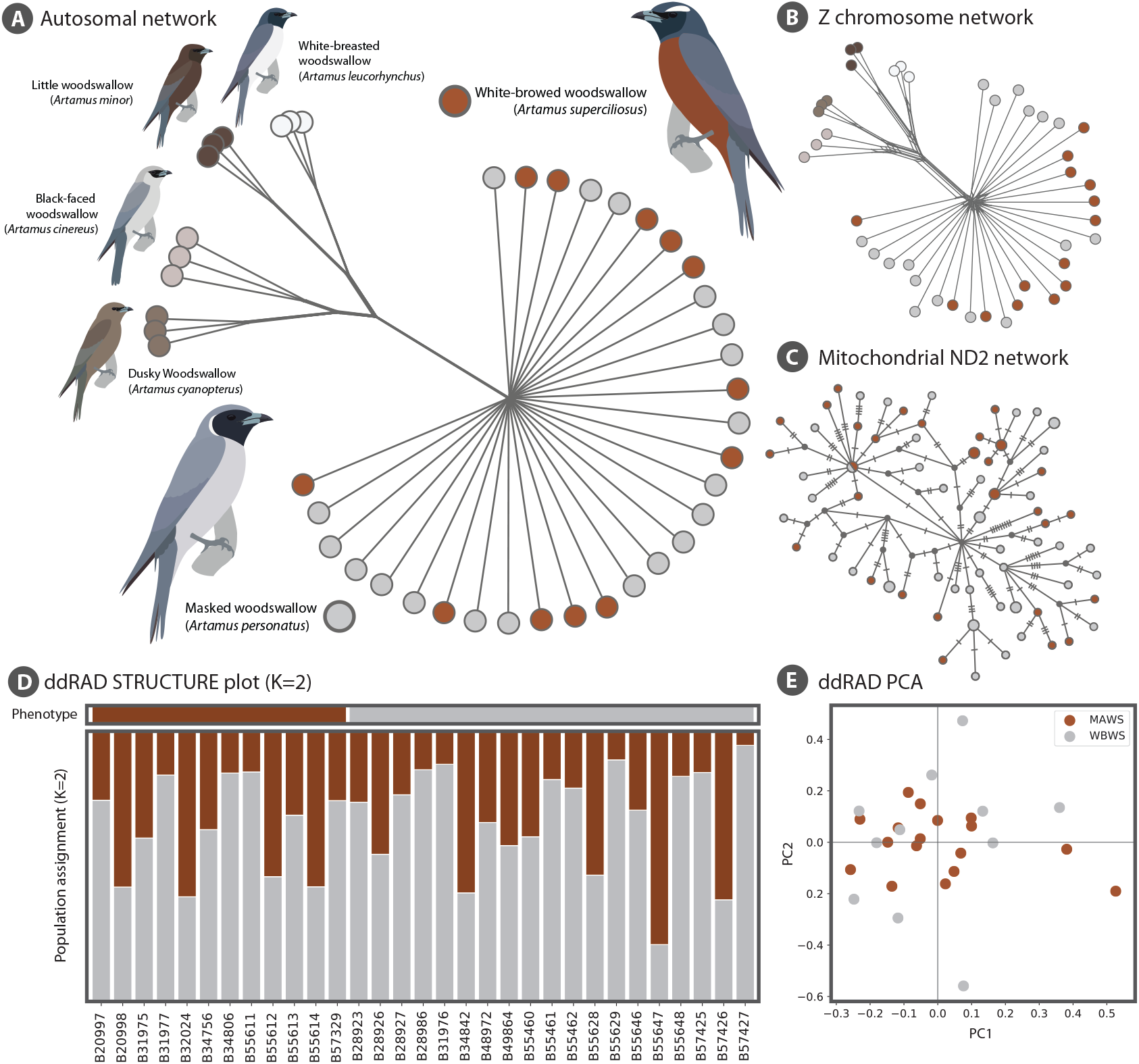
Population structure. The grey and maroon correspond to masked and white-browed woodswallow, respectively. The assignment to species is done purely by plumage. (A) Autosomal network generated from uncorrected pairwise distances among individuals. (B) Z chromosome network generated from uncorrected pairwise distances among individuals. (C) Mitochondrial ND2 haplotype network (note that one internal haplotype is shared due to addition of data since Joseph *et al*. (2006). (D) Population assignment from 20 iterations of STRUCTURE with K=2. (E) Principal components analysis using smartPCA.

To date, mitochondrial DNA is the only available molecular data from the two species. As in the systems cited above, the two species show no discernible population structure in mtDNA despite their striking plumage divergences and they are paraphyletic with respect to each other but together form a sister clade to the rest of *Artamus* (Joseph et al., 2006). The observed mtDNA structure detected in that study is consistent with the later stage of incomplete lineage sorting termed allophyly by Omland et al. (2006; see Figure 1). From this brief summary, we infer that the substantial but incomplete premating isolation has not led to any signal of population structure in incompletely sorted mtDNA. Moreover, we see no clear evidence of the phenomenon of a phenotypic hybrid swarm (Cockayne & Allan, 1926) throughout their extensive range of sympatry in eastern Australia.

In this study, we extend the earlier work on mtDNA (Joseph et al., 2006) to test whether the nuclear genome similarly lacks genetic structure between the two species or whether the previous pattern is a result of extensive mitochondrial introgression. We used ddRAD sequencing to sample loci across the nuclear genome to test whether there is genome-wide divergence across the chromosomes, to describe variation in divergence between autosomes and sex chromosomes, and to test whether we can find peaks of divergence associated with plumage-related genes in our coarse subsampling of the genome. We then inferred the demographic history that could have resulted in the two phenotypic morphs we define today as species. Lastly, we simulated various evolutionary scenarios that could have resulted in the plumage divergence to see which best explains our empirical data.

## Methods

### Sampling

We obtained tissue samples from 19 masked woodswallows (*A. personatus*) and 12 white-browed woodswallows (*A. superciliosus*) from the Australian National Wildlife Collection, CSIRO, Canberra (Table S1). The majority of samples were collected from eastern Australia where the two species are broadly sympatric (supplementary material, Figure S1). Parenthetically, we note that the samples are derived from several decades of opportunistic sampling, reflecting the difficulty of sampling species most frequently seen hundreds of meters into the sky and which have often unpredictable nomadic movements (Higgins, 2006). This means that both species breed opportunistically depending on temperature and rainfall, resulting in different breeding densities across geography and breeding seasons instead of a well-defined breeding range (Joseph et al., 2006; Keast, 1958). We also included three individuals from each of the following species as outgroups: white-breasted woodswallow (*A. leucorhynchus*), little woodswallow (*A. minor*), black-faced woodswallow (*A. cinereus*), and dusky woodswallow (*A. cyanopterus*).

### ddRAD-seq library preparation and bioinformatics processing

We generated DNA sequences using the double-digest restriction-site-associated DNA sequencing (ddRAD-seq) protocol of DaCosta & Sorenson (2014). DNA was extracted using a DNeasy Blood and Tissue Kit (Qiagen, Valencia, CA), and approximately 1 µg of DNA was digested using the restriction enzymes *SbfI* and *EcoRI*. Adapters compatible with Illumina TruSeq reagents and containing unique barcodes were ligated to the digested DNA. Ligated DNA fragments ranging in size from 300 to 450 base pairs (bp) were extracted from 2% low-melt agarose gels and purified using a MinElute gel extraction kit (Qiagen, Valencia, CA). The recovered fragments were amplified using PCR, purified using magnetic AMPure XP beads (Beckman Coulter Inc., Indianapolis, IN), and quantified using real-time PCR with an Illumina library quantification kit (KAPA Biosystems, Wilmington, MA). Uniquely barcoded samples were pooled in equimolar concentrations and sequenced (150 bp reads) on an Illumina HiSeq 2500 at TUCF Genomics, Tufts University (Medford, MA, USA).

Raw Illumina reads were processed using the computational pipeline described by DaCosta and Sorenson (2014) [scripts available at: http://github.com/BU-RAD-seq/ddRAD-seq-Pipeline]. For each sample, identical reads were combined into a single read and recoded with the number of reads and the highest quality score for each nucleotide position. Reads with an average Phred score of <20 were removed. Retained reads from all individuals were clustered into putative loci using USEARCH v. 5 (R. C. Edgar, 2010), with an –id setting of 0.85, and aligned using MUSCLE V. 3 (R. C. Edgar, 2004). Individuals were genotyped at each locus as described in DaCosta and Sorenson (2014): homozygotes were defined when ≥93% of the reads were identical, whereas heterozygotes were defined when a second sequence was represented by ≥29% of reads, or if a second sequence was represented by as few as 10% of reads and the haplotype was confirmed in other individuals. Genotypes were flagged if none of these criteria were met or more than two haplotypes met the criteria. From these flagged genotypes, we retained the allele represented by the majority of reads and scored the second allele as missing data. Similarly, a second allele was scored as missing when the locus was represented by ≤5 reads. We retained all loci that contained ≤10% missing genotypes and ≤5% flagged genotypes.

### Genome placement

We used BLAST to assign the chromosome and location for each ddRAD locus (Altschul et al., 1990). The ddRAD loci were queried against the New Caledonian crow genome (*Corvus moneduloides;* GenBank Accession GCA_009650955.1), the closest relative with a chromosome-scale reference genome available to date. This was done to maximize the number of loci that can be placed along a chromosome and to minimize difference in marker order due to chromosomal rearrangements. To stay consistent with nomenclature used in other studies, we renamed the *C. moneduloides* chromosomes based on the zebra finch nomenclature (Table S2) for our study. The original *C. moneduloides* chromosomes were named by descending order of chromosome length.

### Population structure

The population structure from the nuclear genome was inferred in various ways. First, we used smartPCA from EIGENSOFT (v7.2.1) (http://www.hsph.harvard.edu/alkes-price/software/) to infer the structure through a Principal Components Analysis (PCA). Prior to running the software, the SNPs were filtered to only include variants fixed in the outgroups, non-singletons, and one SNP per ddRAD locus. This was to minimize the effects of incomplete lineage sorting and linkage and to maximize signal. We then ran STRUCTURE using the same set of filtered SNPs (Pritchard et al., 2000). We ran 20 iterations of STRUCTURE for K=1 and K=2 and merged the iterations within K=2 using CLUMPP (Jakobsson & Rosenberg, 2007). The population network was generated using uncorrected pairwise distances among individuals. The network for the autosomal and Z-linked loci were inferred separately. Unlike for the PCA, the SNPs were not filtered for this analysis. The population network was then visualized using SplitsTree (Huson, 1998). Finally, the mitochondrial ND2 haplotype network was inferred and visualized using the TCS network implemented in PopArt (Leigh & Bryant, 2015). The ND2 sequences were downloaded from GenBank (accession numbers in Table S1).

### Population genetic parameters

All population genetic parameters were calculated using custom scripts and the following equations. Population differentiation was estimated using Hudson’s F_ST_ which is less influenced by differences in effective population size (Bhatia et al., 2013; Hudson et al.,1992). We used the following equation: 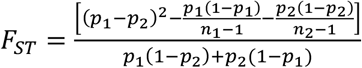, with *p* and *p* being the frequency of the allele in population 1 and population 2, respectively, and n_1_ and n_2_ being the number of chromosomes sampled in each population. Average F_ST_ was calculated within each ddRAD locus and across the genome using the ratio of averages. Absolute population divergence (D_XY_) was estimated using the following equation 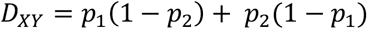 divided by the total length in base pairs. Genetic diversity within each species was calculated using both Tajima’s (π) and Watterson’s estimator (θ_W_) using the equations 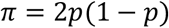and 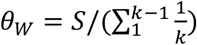 divided by the total length in base pairs with *p* being the allele frequency of either allele, S the number of segregating sites and *k* the number of chromosomes. Lastly, Tajima’s D was calculated using 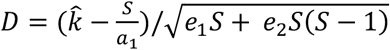. The definitions for the different variables can be found in Tajima (1989).

### Demographic modeling

We used fastsimcoal2 (Excoffier et al., 2021) to test various demographic scenarios between *A. superciliosus* and *A. personatus* using the unfolded 2D joint site frequency spectrum (SFS). The SNPs were filtered to retain only those fixed for all three individuals in all four outgroup species and we subsampled a single SNP per ddRAD locus to allow for an unfolded joint frequency spectrum of unlinked sites. Using only SNPs that are fixed across all outgroup individuals would minimize the likelihood of mis-polarization by incorrectly classifying a segregating allele as ancestral. We assumed the allele fixed for all outgroup species to be the ancestral allele. We also omitted sex-linked loci and outlier loci to minimize effects of differences in effective population size and strong selection within the genome. This retained 4169 unlinked, autosomal SNPs. We tested models of panmixia, recent divergence, isolation-with-migration, secondary contact, and change in migration. In order to be comparable to the other models, panmixia was simulated as two populations with a migration rate m = 0.5 which corresponds to full sympatry (Coyne & Orr, 2004). The change in migration scenario includes two time periods with differing levels of migration to reflect a history of low migration during divergence followed by a period of higher migration which homogenized the divergence. We created bootstrapped SFS by choosing a single random SNP within each ddRAD locus so the model selection is not influenced by stochastic differences in SNP selection. For model selection, we ran each model 100 times for 10 bootstrapped SFS and the run with the best likelihood for SFS was chosen.

When performing parameter estimation after choosing the best model, we discovered two optimal parameter combinations where either masked or white-browed woodswallow would have the larger effective population size and corresponding migration rate. To differentiate between these scenarios, we performed another test comparing a model where migration rate parameter space is bound such that: *m_MAWS®WBWS_* > *m_WBWS®MAWS_* versus *m- _WBWS®MAWS_* < *m_MAWS®WBWS_.* We fixed parameters that had low variance from our initial runs and let other parameters such as effective population size still vary. We then chose the scenario with the higher likelihood and restricted the migration rates while estimating the variance of the parameters. Due to this lack of convergence when estimating parameters for the best model, we also estimated parameters under the two next best models to see how well they would perform.

To obtain the variance in the parameters, we bootstrapped the SNPs to generate 50 SFS and reran the parameter inference 50 times for each. The parameters that were previously fixed were allowed to vary for this final parameter estimate. The best run for each SFS was chosen and the median and variance (median absolute deviation) was calculated across the 50 bootstrapped SFS.

### F_ST_ outlier analysis and gene ontology search

Because both species are largely sympatric throughout the range of white-browed woodswallow and the sampling was mostly done in the same locations (Figure S1), we did not need to test whether the F_ST_ outliers are due to isolation-by-distance as you might in a classic parapatric scenario. Each F_ST_ value corresponds to a single ddRAD locus rather than a single SNP. We used a 99.5 percentile threshold to identify putative outliers. Of these outliers, we also investigated four loci which were substantially more differentiated than the rest. We then identified which genes were 40kbp upstream and downstream of these outliers using the genome annotation of *Corvus moneduloides*. We retrieved the names and molecular functions of these genes based on homology to the chicken (*Gallus gallus*) annotation. We also searched whether we had ddRAD loci within the following known plumage genes from other studies: *ASIP, FST, CYPJ19, MC1R, TYR, TYRP1, OCA2, SLC45A, KIT, DCT,* and *Mitf* (Funk & Taylor, 2019; Kim et al., 2019; San-Jose et al., 2017; Toews et al., 2016; Toomey et al., 2018; Twyman et al., 2018).

### SLiM simulations

To investigate the potential underlying mechanism that resulted in the outlier loci, we simulated various evolutionary scenarios that could result in high F_ST_. Using SLiM3 (Haller & Messer, 2019) we simulated a 500bp sequence to represent an independent ddRAD locus under different scenarios. We simulated five main scenarios: *neutral, background selection, balancing selection, ancient sweep,* and *recent sweep*. For *neutral, background selection,* and *balancing selection* we also had a scenario with and without gene flow to reflect freely moving and barrier loci, respectively. Within *ancient sweep* and *recent sweep*, we also tested combinations of *hard sweeps* vs. *soft sweeps*. This resulted in a total of 12 different scenarios. The *neutral* and *background selection* were our null models where divergence was not driven by differences in selection regimes (Comeron, 2017). Both mutation and recombination rates were kept constant and we used the parameters estimated by the demographic modeling for population sizes, divergence times, and gene flow. We simulated each scenario 100 times and extracted the F_ST_ and D_XY_ between populations and the θ and π within populations. Specific details of the simulations can be found in the supplementary methods.

## Results

### ddRAD sequencing

We obtained an average of 869 568 reads per individual (median = 754 301; range = 351 718–4 669 735). After filtering and assembly, we recovered a total of 4664 ddRAD loci containing 76 336 SNPs in 582 775 bp. Of these, 50 913 SNPs in 4371 loci were fixed in all four outgroup species. We were able to locate the genomic position of 4428 loci in the New Caledonian Crow assembly (181 of which mapped to the Z chromosome).

### Population structure, differentiation and genetic diversity

We found no population structure in the nuclear ddRAD loci between the two species (Figure 1A to E). Both the autosomal and Z-linked markers (Figure 1 A, B) showed a distinctive star-like pattern indicative of a fairly panmictic population despite divergence in many plumage traits. The STRUCTURE plot for K=2 (Figure 1D) also shows no discernible population structure. The mean Ln likelihood of the data across the 20 runs is -63104.35 (± 23.99) for K=1 and -78574.17 (±13503.70) for K=2 suggesting a model of no population structure fits better. The PCA (Figure 1E) also does not separate the two species into distinct clusters.

Consistent with the lack of population structure, the genetic differentiation between the two species is very low, the average autosomal and Z chromosome F_ST_ being 0.003 and 0.0002, respectively. Although the average F_ST_ is an order of magnitude lower in the Z chromosome and contrary to expectation from differences in effective population size between marker types, the median F_ST_ is 0.0 for both autosomes and Z chromosome. This suggests that the elevated F_ST_ in the autosomes is driven by the few outliers (Figure 2). The global D_XY_ between the two species is D_XY_ = 0.012 per bp. The estimates for genetic diversity within the species were similar: π_WBWS_=0.0118, π_MAWS_=0.0120, θ_WBWS_ = 0.0176, θ _MAWS_=0.0199. The estimates of Tajima’s D within each species were D_WBWS_= -1.026 and D_MAWS_= -1.184. The comparisons among parameters of the per locus estimates can be found in Figure 2.

**Figure 2.**
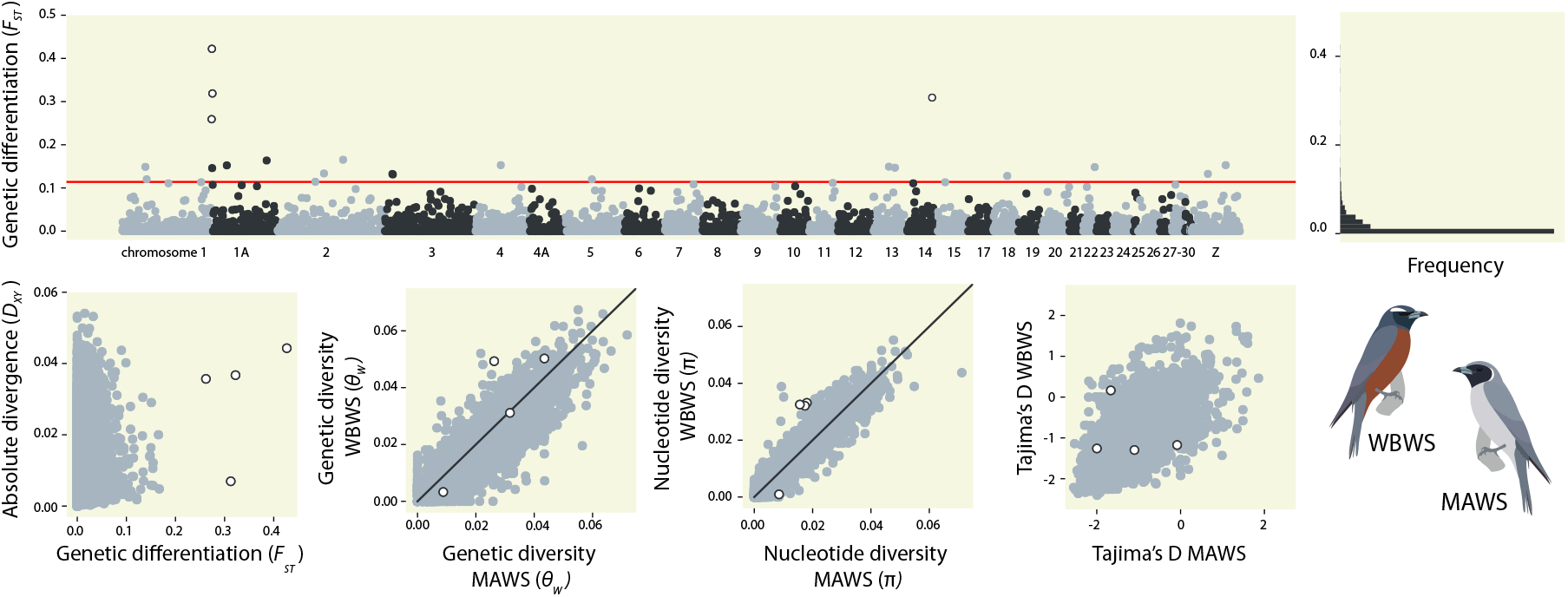
Genome differentiation and outlier loci. Top left: Genetic differentiation across the ddRAD loci by physical order on the *C. moneduloides* reference genome. The alternating colors represent different chromosomes as numbered below. The red line corresponds to the 0.995 percentile threshold. The open circles represent the extreme outlier loci. Top right: Histogram of estimates of genetic differentiation across ddRAD loci. Bottom (from left to right): Comparison between absolute and relative genetic divergence among ddRAD loci, comparison of genetic diversity (Watterson’s theta) among the ddRAD loci between the white-browed woodswallow (WBWS) and masked woodswallows (MAWS), comparison of nucleotide diversity (π) among the ddRAD loci between the white-browed and masked woodswallows, and comparison of Tajima’s D among ddRAD loci between the white-browed and masked woodswallows

### Demographic modeling

Five different demographic scenarios were tested. Of the five, the change in migration scenario had the best support (Figure 3). Secondary contact (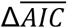 = 107.27) and isolation-with-migration (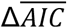 = 118.09) had slightly lower support. The scenarios of recent divergence (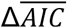 = 840.98) and panmixia (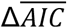 = 3174.69) had the lowest support. For the change in migration scenario, the estimations of migration rates were independent from one another so the models were allowed to explore scenarios where initial migration was higher than contemporary migration. Only models where contemporary gene flow was higher than ancient gene flow were recovered in our iterations.

**Figure 3.**
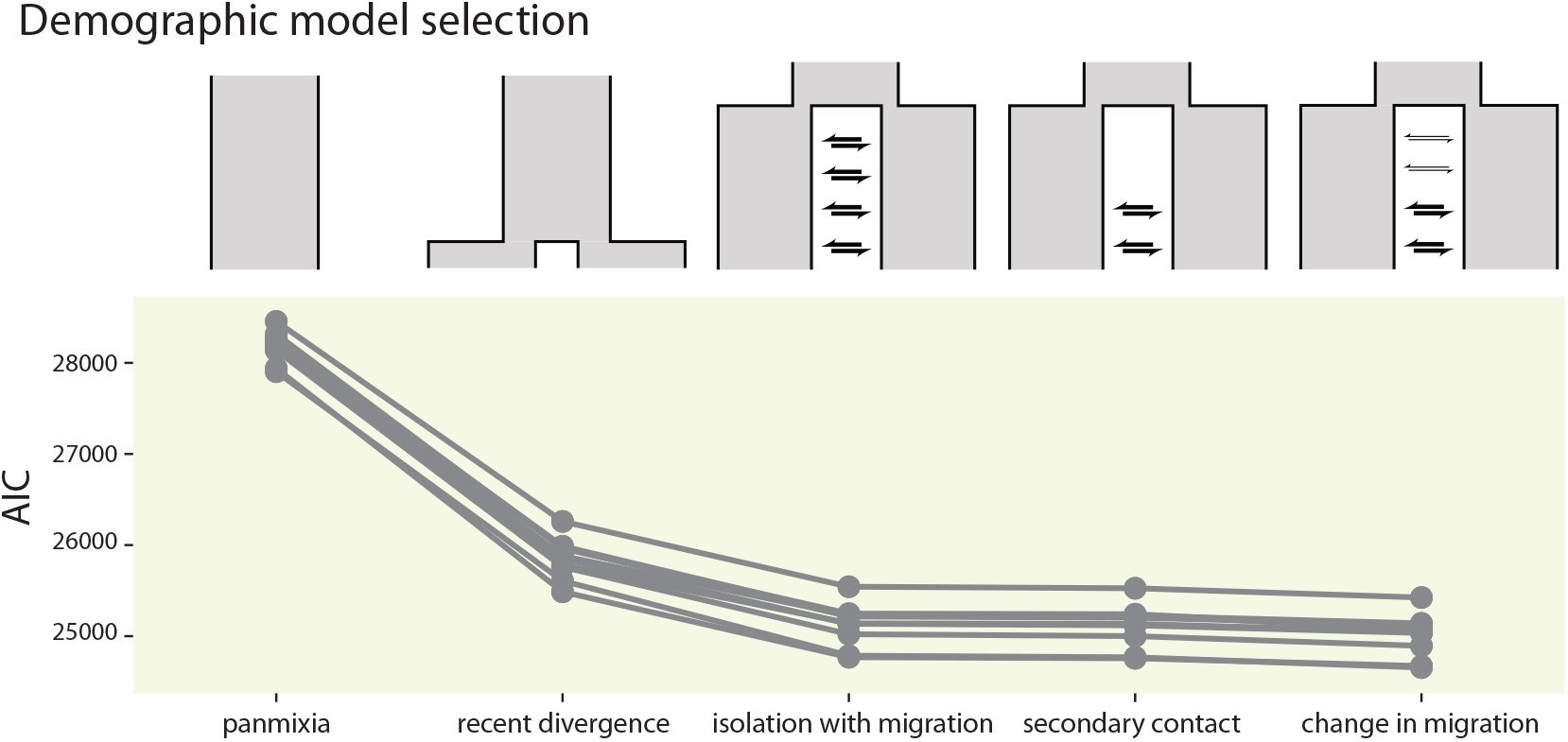
Demographic modeling. Demographic inference performed using fastsimcoal2. *C*omparison of AIC across different demographic scenarios among 10 bootstrap replicates of the SFS.

The change in migration model consistently had higher likelihood in all ten bootstrap replicates but the parameter estimates had two different optima. That is, either the masked or white-browed had a substantially larger effective population size and gene flow relative to the other. The resulting estimates for effective population sizes of the masked woodswallow were 20-fold more than white-browed (Table 1) which is contradictory to the similar levels of genetic diversity. This may be due to overfitting and lack of signal in the data. Therefore, we report the parameter estimates of the second-best model: secondary contact. The estimates of the effective population sizes were more similar between the masked and white-browed woodswallows, consistent with the estimates of genetic diversity. Parameter estimation suggests that the ancestral population size was 2N = 252 887, and that of masked and white-browed woodswallows were 2N = 1 325 798 and 1 159 530 individuals, respectively. This suggests that the both the masked and white-browed woodswallows experienced an expansion from their ancestral population size. Contemporary estimates of gene flow values were 10.85 and 94.37 migrants/generation, from masked to white-browed and from white-browed to masked, respectively. The median estimated time since divergence was 195 256 generations (± 136 445 generations) and median estimated time of secondary contact was 13 079 generations (± 5 801 generations). These convert to 347 554 and 23 280 years, respectively, assuming a generation time of 1.78 years (Higgins, 2006).

**Table 1.** Demographic parameter estimation. Parameter estimation under the three best models simulated in fastsimcoal2. These are the median and variance (median of absolute deviations) of best parameter estimates from 50 iterations of 50 bootstrapped SFS.

| Model |  |  | Isolation with migration | Secondary contact | Change in migration |
| --- | --- | --- | --- | --- | --- |
| Population size<br>(2N) |  | Ancestral | 93 454 ± 62 077 | 252 887 ± 175 643 | 16 834 ± 16 668 |
|  |  | MAWS | 529 013 ± 296 300 | 1 325 798 ± 623 541 | 1 222 182 ± 473 361 |
|  |  | WBWS | 288 910 ± 183 225 | 1 159 530 ± 751 609 | 59 250 ± 15 829 |
| Gene flow<br>(2N) | Contemporary | MAWS → WBWS | 7.06 ± 4.73 | 10.85 ± 7.20 | 11.19 ± 7.46 |
|  |  | MAWS ← WBWS | 52.72 ± 9.73 | 94.37 ± 31.35 | 58.53 ± 7.17 |
|  | Ancient | MAWS → WBWS | 7.06 ± 4.73 | 0.00 | 32.33 ± 3.27 |
|  |  | MAWS ← WBWS | 52.72 ± 9.73 | 0.00 | 0.15 ± 0.06 |
| Time<br>(generations from<br>present) | Divergence |  | 42 288 ± 26 395 | 195 256 ± 136 445 | 604 406 ± 129 955 |
|  | Secondary contact |  | - | 13 079 ± 5 801 | - |
|  | Change in migration |  | - | - | 584 845 ± 118 704 |

### Outlier loci

Based on the 99.5 percentile cut-off we recovered 22 outlier loci. Of these 22, four had substantially higher F_ST_ than the others. Three of these loci were located at the distal end of chromosome 1A and the fourth was on chromosome 14. The three loci in chr1A were also adjacent to two other outliers above the 99.5 percentile cut off. This region encompasses at least 700kb at the distal end of the chr1A. The loci in chr1A also had high D_XY_ (Figure 4) suggesting they are potentially barrier loci whereas the locus in chr14 had low D_XY_, which might suggest background selection or a selective sweep. The genetic network generated for the 1.5 Mb region of chr1A (Figure 4) suggests higher derived diversity in the white-browed woodswallow and retained ancestral polymorphism for the masked woodswallow.

**Figure 4.**
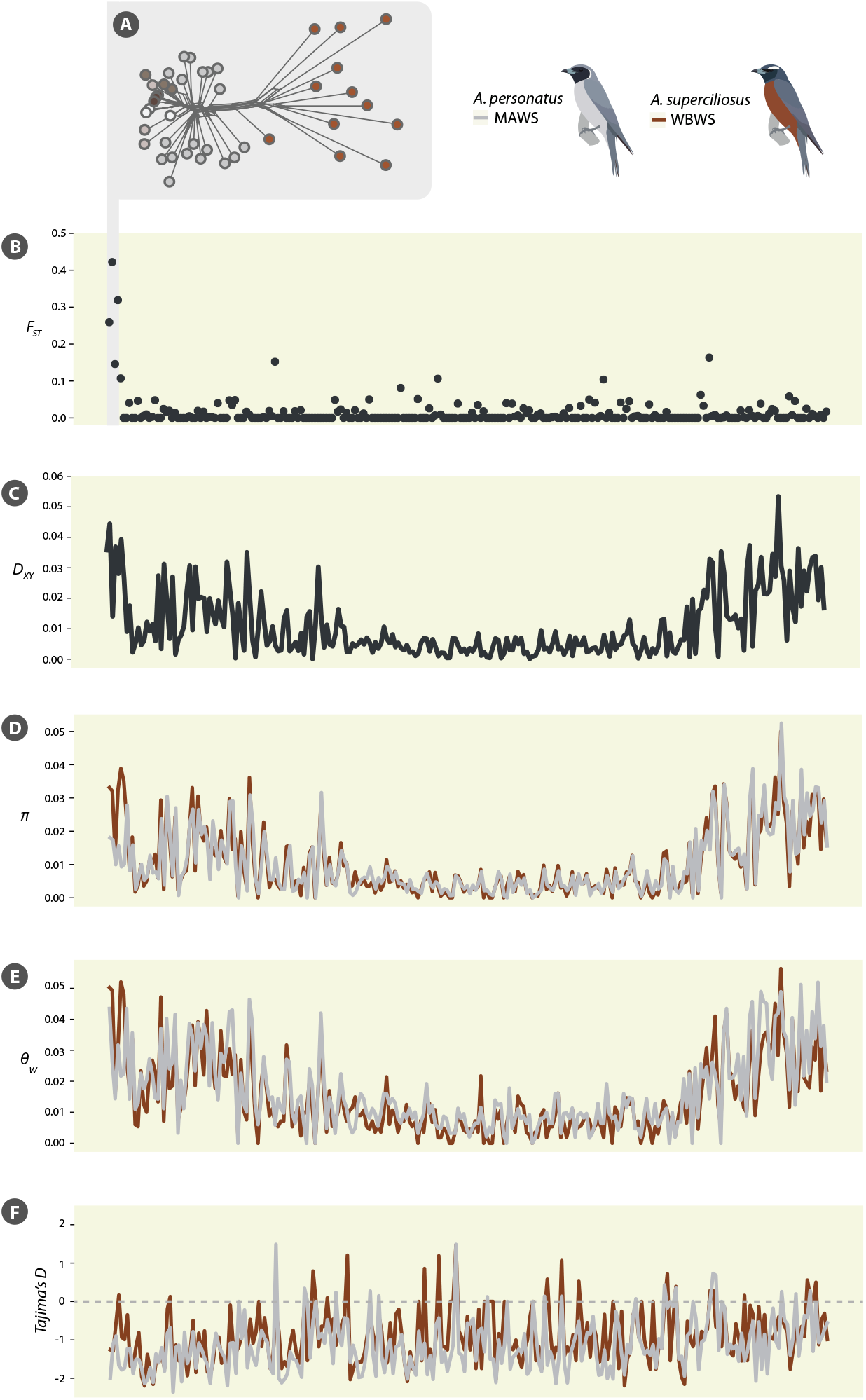
Differentiation and diversity along chromosome 1A. (A) Uncorrected genetic distance network of from the 5 outlier loci suggesting masked woodswallow has retained ancestral variants from the outgroups and white-browed woodswallow is likely the derived form. (B) Relative genetic differentiation across the length of chromosome 1A. (C) Absolute genetic divergence (D_XY_) across the length of chr1A suggesting both elevated relative and absolute measures of divergence. (D) Nucleotide diversity (π) across the length of chr1A suggesting higher diversity in the white-browed woodswallow. (E) Genetic diversity (θ) across the length of chr1A suggesting slightly higher diversity in the white-browed woodswallow. (F) Tajima’s D across the length of chr1A showing overall negative values but not substantial difference between the two species.

There was a total of 36 genes within 40kb on either side of the 22 outlier loci (Table S3). Of the outlier loci with the highest differentiation, only one (*SOX5*) is found within or adjacent to the 700kb in chr1A and two (*AXIN1* and *PDIA2*) are found adjacent to the chr14 outlier locus. The four outlier loci in chr1A span 696kb-1.36Mb while *SOX5* spans 737kb- 1.2Mb, and the outlier locus in chr14 falls directly within the *AXIN1* gene coordinate range. Both *SOX5* and *Axin1* are involved in negative regulation of the *Wnt/*/β-catenin signaling pathway, which regulates feather development (Chang et al., 2004; Xie et al., 2020). Of the previously characterized plumage genes, only one had an associated ddRAD locus and it did not have high differentiation between species.

### SLiM simulations

Our simulations suggest that only very little neutral differentiation can occur given the time of divergence and large effective population sizes of the two species (Figure 5; Figure S10). This is consistent with the observed overall lack of genome-wide differentiation. All other scenarios, on the other hand, can generate elevated F_ST_ and D_XY_ relative to the neutral loci. Loci under *balancing selection* (with migration) and *hard sweeps* (both ancient or recent) were qualitatively most similar to our outlier loci (Figure 5). The other empirical observation we used for comparison is the larger genetic (θ) and nucleotide diversity (π) in white-browed woodswallows relative to masked in the outlier loci (Figure 2, Figure 4D & E). The only scenarios under *balancing selection* where this is true was in loci with gene flow; this would be inconsistent with the population structure in these loci (Figure 4A). The only remaining scenarios that are consistent with all empirical patterns are *hard sweeps* that occurred either when the populations diverged or after they came into secondary contact. The patterns between these two scenarios are qualitatively similar. This is likely due to the very recent divergence time estimate. None of the simulations reached a full sweep and that is likely due to the short times and large effective population sizes. Only very large selection coefficients, which are even less likely in nature, could have likely resulted in a complete sweep. We were only able to make qualitative comparisons between empirical data and simulations based on relative rather than absolute values. This is because we did not have information on many parameters (mutation rate, recombination rate, selection coefficient, etc.) that would result in deviation of the absolute values and quantitative differences.

**Figure 5.**
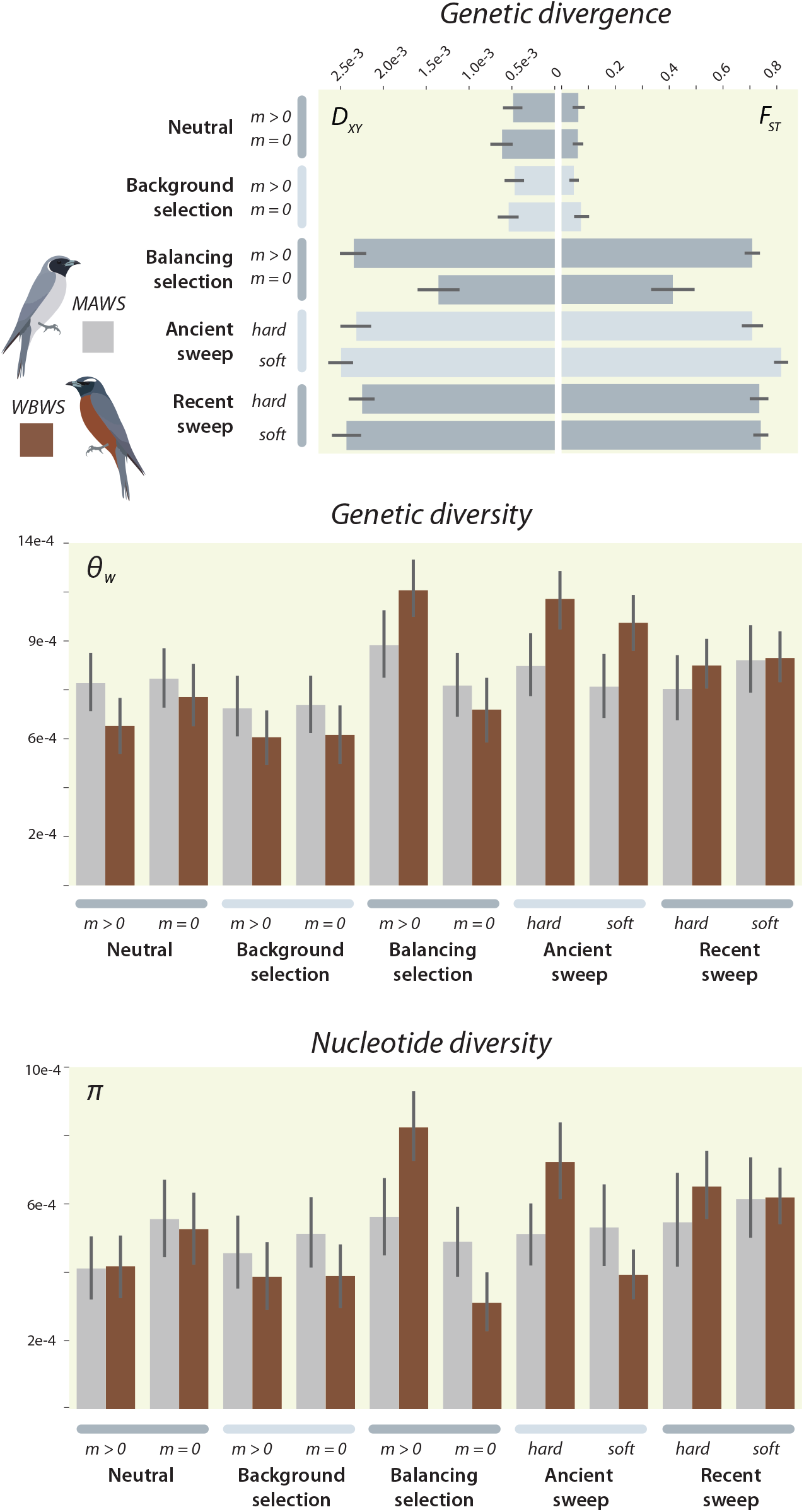
Simulations. Evolutionary scenarios simulated using SLiM3 to test potential origin and maintenance of the outlier loci. Each scenario was run 100 times and the bars reflect the standard deviation of the estimates among the 100 iterations. MAWS – masked woodswallow; WBWS – white-browed woodswallow.

## Discussion

Divergence in secondary sexual traits such as plumage is often associated with isolation via sexual selection. Counter examples are accruing in avian systems where plumage divergence is maintained despite genome-wide homogenization (e.g., Knief et al. 2019; Toews et al. 2016). Such systems often involve geographically localized hybridization or introgression that effectively homogenizes most of the genome. Here we studied Australia’s masked and white-browed woodswallows *Artamus* spp in their broad sympatry over half of the Australian continent and between which phenotypic hybrids, though not unknown, are rare. We investigated the genomic basis to their high phenotypic divergence, which is sustained despite a lack of genetic divergence.

Our study using nuclear ddRAD markers further supports the lack of structure and divergence found in the mitochondrial DNA (Joseph et al., 2006). We used various methods to infer population structure and so capitalize on different attributes of the nuclear DNA data, yet still we found little evidence of allele frequency differences. The population network of the autosomal data, for example, shows a star-like pattern among the individuals of both species far more clearly than that of mtDNA (Figure 1). Next, the mtDNA data might suggest either no divergence has occurred or that the mitochondrial genome from one species has ‘swamped’ or replaced the other. The lack of shared haplotypes, however, is unexpected under a simple mtDNA capture. Although we still lack sampling of the masked woodswallow from the western half of the continent where that species is prevalent and so may have missed within-species structure or unique alleles, a key new finding is that the nuclear genomes of the two species are largely indistinguishable throughout their sampled and nonetheless vast range (autosomal F_ST_ = 0.002, Z chromosome F_ST_ = 0.0003, Figure 2).

Despite being the only two species in *Artamus* exhibiting sexual dichromatism (Schodde & Mason, 1999), there is no elevated differentiation in the Z chromosome. This contrasts with other avian systems having high plumage differentiation coupled with low autosomal and high Z genetic differentiation (Dhami et al., 2016); the latter “large Z effect” is especially prevalent when adaptive differences are involved (Ellegren, 2009; Schield et al., 2021). Mank (2009) noted the complex connections between sex chromosomes and sexual dimorphism, however, and called for tests of assumptions such as we have presented here. The largely nomadic and in some years, strongly migratory, nature of these woodswallows likely allows for sympatry without strong competitive exclusion. This broad sympatry also means that occasional hybridization is not geographically restricted and likely happens throughout much of south-eastern Australia where the birds mostly are known to breed. This would facilitate introgression and genome homogenization across both species and concomitantly hinder the accumulation of genome-wide divergence.

Despite the observed lack of population structure, our demographic analyses have poor support for panmixia or recent divergence. Although the demographic scenario with the best support was that of a change in migration rate during divergence, the scenario of secondary contact resulted in more reasonable parameter estimates which were also consistent with the estimated population genetic parameters. The inferred recent divergence times, contemporary gene flow, and large population sizes are likely why we do not detect population structure. This is further corroborated by our simulations (see below). Our inference suggests that the two species experienced secondary contact ∼23 kya, roughly around the last glacial maximum (LGM), where one species likely expanded into the other’s range. This scenario is not unlike another Australian arid-adapted species, the butcherbirds *Cracticus* spp, which experienced range expansion and secondary contact during the LGM (Kearns et al., 2014). The higher migration rate from white-browed into masked woodswallow is consistent with a scenario where the masked woodswallow invaded the white-browed’s range thereby retaining more of the native species’ alleles in the invading species’ genome (Joseph et al., 2006; McElroy et al., 2020; Rheindt & Edwards, 2011). This contrasts with the scenario where alleles of the population with the lower effective population size (i.e., white-browed woodswallow) are purged due to high deleterious load (Moran et al., 2021) as both species have similarly large population sizes. The derived plumage morph then likely established rapidly and is evidently resistant to gene flow after the secondary contact. We acknowledge that the absolute parameter estimates should be interpreted with caution. We tested 2-population models based on plumage rather than genetically defined demes. Rather, we emphasize the relative values not absolute point estimates.

Extensive homogenization of genomes can reveal regions that may be involved in phenotypic divergence and speciation, and by extension the underlying evolutionary processes, which are often masked by a ‘noisy’ landscape of divergence. Under the inferred demographic history, we simulated different evolutionary processes that could have led to the pattern of outlier loci we observed. Our simulations suggest that very little neutral differentiation would occur even in strict allopatry as the divergence times are too recent and population sizes too large for drift to work efficiently. The elevated F_ST_ and D_XY_ are then likely due to the outliers in chr1A acting as barrier loci with restricted gene flow. The simulations that resulted in the elevated genetic diversity similar to the chr1A outliers are hard selective sweeps occurring either after divergence or at secondary contact, and possibly any point in between. In contrast to chr1A, the outlier in chr14 has elevated F_ST_ but low D_XY_, which can result from long-term background selection resulting in reduced genetic diversity (see Cruickshank & Hahn, 2014). Our simulations under different levels of background selection, however, did not result in low D_XY_ and high F_ST_ relative to neutral loci as predicted (Figure S10). A denser sampling of SNPs adjacent to the outliers would provide better signal from linkage to help distinguish between these different scenarios.

Importantly, we can begin to formulate hypotheses on the relevance of the genes associated with the outlier loci. Here, we focus on *SOX5* in chr1A and *Axin1* in chr14 where the extreme outlier loci fall within the boundaries of the respective genes. Neither of these genes have been implicated in previous studies of plumage divergence but both are involved in negative regulation of the *Wnt/β-catenin* pathway, which is well-known in regulating feather development (Chang et al., 2004; Xie et al., 2020). There is no documented direct interaction between *SOX5* and *Axin1*. *SOX5*, however, induces the production of *Axin2*, which is a negative regulator of the *Wnt*/β-catenin signaling pathway and functionally equivalent to *Axin1* (Dao et al., 2007; Quiroga et al., 2015). Overexpression of *Axin1* and *Axin2* reduces β-catenin levels and signaling. Another gene, *G3BP1*, found near an outlier locus in chr13 is also a negative regulator of the *Wnt*/β-catenin signaling pathway. *SOX5* additionally has been associated with strong phenotypic effects such as the pea-comb phenotype in chickens (Wright et al., 2009) and pigmentation through melanocyte development in mice and medaka fish (Nagao et al., 2018; Stolt et al., 2008). More precise delineation of boundaries of highly differentiated genomic regions and other potential candidate genes associated with phenotypic divergence requires denser sampling of SNPs, preferably through whole genome resequencing.

### Future prospects

Having established that the nuclear genomes of white-browed and masked woodswallows show little divergence and no population structure, this species complex offers multiple avenues for investigation of the genetic architecture and maintenance of plumage divergence. Assembling and annotating a high-quality reference genome for the two species would place the variants in the correct physical order and may even reveal unique structural variation between the species. Furthermore, population-level whole genome resequencing would provide a more complete sampling of SNPs across the genome. This can validate the outliers presented in this study and potentially identify other candidate genes missed by the ddRAD sequencing. Yet, some causative small-effect loci involved in this complex trait may remain hidden despite the homogenizing effect of gene flow because of the high genetic variation within each species (Le Corre & Kremer, 2003; Yeaman, 2015). Additional sampling of masked woodswallow individuals from the part of their range where they do not overlap with the white-browed woodswallow may reveal lineage specific alleles that have not introgressed. This would be logistically non-trivial given both the remoteness and vastness of their range in western parts of Australia, the often-nomadic movements of the birds and indeed that they are often seen high in the sky prohibiting sampling. Conversely, masked woodswallows could more readily be sampled if encountered when a flock can be repeatedly located over several days feeding in an area of low, flowering shrubs (e.g., Higgins 2006). Both species are increasingly kept in captivity where gene expression studies during feather development could test whether the genes found in this study play a direct role or interact with other candidate genes. Field studies on mate choice and sampling hybrids, again difficult logistically due to the birds’ often unpredictable occurrences, would nonetheless disentangle the role of premating and postzygotic isolation in maintaining plumage divergence despite extensive introgression.

Systems such as we have examined here with striking phenotypic divergence yet little genomic divergence contrast with those having little to no discernible phenotypic divergence and high genomic divergence (i.e., “cryptic species” (Fišer et al., 2018)). Although these systems may be the exceptions to the rule, they bring to light the complexity of understanding speciation and species delimitation based on these metrics of divergence. Our study adds to those exemplifying divergence in only a few loci strongly influencing divergence of a secondary sexual trait such as plumage. We also show two further key findings: strong plumage divergence does not necessarily prevent whole-genome introgression and genes underlying a secondary sexual trait such as plumage are more likely to be found in autosomes rather than sex chromosomes. As we build a better understanding of the genetic architecture and evolution of these individual traits, we can piece together a more complete picture of the origin and evolution of biodiversity.

## Acknowledgments

Specimens used in this study were collected under all appropriate ethics and scientific collecting permits over several decades and with the help of many landholders as acknowledged in an earlier paper (Joseph et al. 2006). For the present study, we would like to thank Sara Seibert for assisting with laboratory work and Christine Darwood, Graeme Chapman, Bastiaan Hensen, Mick Roderick and Andrew Silcocks for kindly granting permission to use their photographs in the Supplementary Material.

## Data accessibility

Raw Illumina reads are deposited in GenBank Accession [TBD] Processed ddRAD fasta files are available in DRYAD doi [TBD].

## Author Contributions

J.V.P analyzed the data and wrote the manuscript. J.L.P. performed the lab work, processed and analyzed the data, and edited the manuscript. L.J. designed the research, and helped write and edit the manuscript.

## Notes

### Competing Interest Statement

The authors have declared no competing interest.

### Summary of Updates

New parameter estimates were performed and the simulations redone. The tables in the Supplementary Material have been revised to include additional information on sample numbers.

