## Supplemental Material for "Sustained plumage divergence despite weak genomic differentiation and broad sympatry in sister species of Australian woodswallows (*Artamus* spp.)"

**Supplemental Information for:**

### Sustained plumage divergence despite weak genomic differentiation and broad sympatry in the sister species masked- and white-browed woodswallows

Joshua V. Peñalba, Jeffrey L. Peters, Leo Joseph

#### Supplementary methods

##### *SLiM simulations*

We used SLiM3 (Haller and Messer 2019) to compare potential underlying processes that may have driven the differentiation of the outlier loci in the woodswallow system. We tested five main processes (explained in detail below) with variations within: *neutral*, *background selection*, *balancing selection, ancient sweep*, and *recent sweep*. Each simulation is an independent 500bp fragment meant to represent unlinked ddRAD loci evolving under the different scenarios in the genome. For all simulations, we fixed mutation rate to be $7 \times 10^{-10}$ and recombination rate to be $1.4 \times 10^{-9}$ based on the zebra finch estimates (Singhal et al. 2015). Although this would likely not be an accurate estimate of the mutation and recombination rate in the woodswallows, because we are only using simulation for qualitative comparisons changing the mutation and recombination rate will change the estimates of specific parameters but not the relative differences they have among each other. These simulations assume a fairly uniform mutation and recombination rate within the genome as we are not testing whether this difference could be driven by large variation in within-genome mutation and recombination rate. The simulations are also meant to reflect the inferred demographic history so we set the ancestral population to 125,000 and the daughter populations to 660,000 and 580,000 for the masked- and white-browed woodswallows, respectively

In order to reduce the time it takes to perform the simulations, we rescaled all factors by 100. This means that the simulated populations and the timing of events are 100x smaller than the empirical estimates. This also means that the rates (mutation, and recombination) are 100x larger to compensate. The migration rates stay the same as it would decrease to adjust for the smaller population size and increase by the same factor to adjust for the shorter time. From this point forward, we will be referring to the rescaled values. The ancestral population was allowed to accumulate mutations and reach equilibrium by letting it evolve for 20,000 generations (more than 10x the effective population size of 1250 as recommended by the software developers). The daughter populations were then split and evolved without gene flow. The secondary contact was set at generation 21,822 with migration from white-browed to masked woodswallow set to 1.5 x $10^{-4}$ and the inverse set to 5.5 x $10^{-6}$. For simulations where the locus serves as a barrier to gene flow, the migration rates were set to 0. Finally, the populations were sampled at generation 21,952 which reflects current time as inferred by the demographic models. At this time-point, 40 and 28 chromosomes, corresponding to 20 and 14 diploid individuals were sampled from the masked- and white-browed woodswallow populations respectively. This reflects the empirical sampling we had in this manuscript.

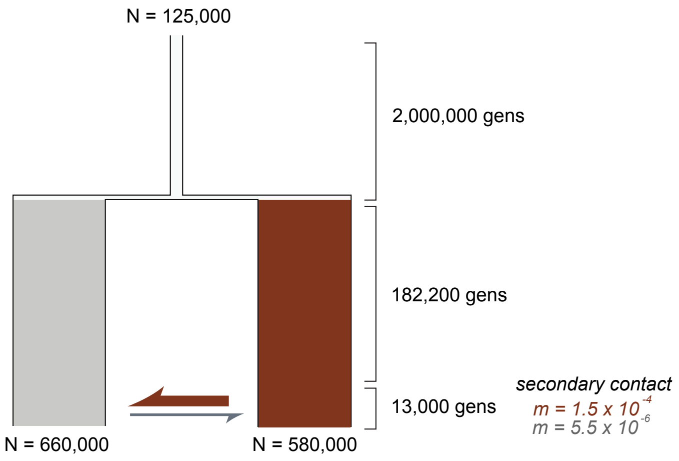


*Simulation schematic*
Values of the population genetic sizes, generation times, and gene flow before rescaling. The numbers are not identical to the estimates from the demographic model but adjusted for convenience. The diagram is not proportional.

**NULL SCENARIOS**

*Neutral (with gene flow, without gene flow)*

The neutral scenario is the basic null scenario with no type of selection is induced at any point in the history of the populations. The only difference is whether secondary contact and gene flow occurs at generation 21,822 or the loci remain as barrier loci with gene flow at 0.

*Background selection (with gene flow, without gene flow)*

The background selection scenario is an alternative null scenario where the whole region experiences an excess of mutations with a negative fitness effect. This is meant to keep diversity low relative to a fully neutral scenario. This reduction in within population diversity can result in peaks in differentiation (Cruickshank and Hahn 2014). This scenario is still considered a null hypothesis as this selection is uniform among populations rather than a differential form of selection that may drive speciation (Comeron 2017). To simulate background selection for the entire 500bp locus, 80% of mutations were neutral while 20% of mutations were under negative selection. The mutations under negative selection receives a fitness effect sampled from a gamma distribution with a mean of -0.05 and a shape parameter (alpha) of 0.2. We also tested different increasing strengths of background selection (-0.1, -0.2, -0.4, and -0.6) to see how stronger background selection would affect the differentiation of the loci.

**DIFFERENTIAL SELECTION SCENARIOS**

Because the derived diversity seems to have occurred in the white-browed woodswallow (Figure 4A), the sweep mutation under the following scenarios occurred in the white-browed woodswallow after it split from the masked. Although this mutation has varying selection regimes in the white-browed woodswallow, it is set to be selectively neutral in the masked woodswallow. The mutation is set to occur immediately after population split in the middle of the locus (250bp) except in recent sweep scenarios.

*Balancing selection (with gene flow, without gene flow)*

Balancing selection in our simulations refer to the maintenance of a polymorphism by the derived being under positive selection in lower frequencies but shifts to negative selection in higher frequencies. This minimizes the chance of the allele being lost or fixed in the population and the constant shift in selection regime results in higher genetic diversity in the population where the balancing selection is occurring. We tested this scenario because of the high genetic diversity in the white-browed woodswallow associated with the outlier locus in chr1A. We decided a threshold of high allele frequency by sampling a random number between 0.75 and 0.95. If the derived allele exceeds this frequency, the selection coefficient shifts to -0.2 and if the derived allele goes back below this frequency the selection coefficient shifts back to 0.5.

*Ancient sweep (hard, soft)*

Under ***hard sweeps***, the mutation is under positive selection the moment it occurs. Under ***soft sweeps***, the mutation is neutral when it occurs and is not beneficial until it reaches a certain threshold. One can think of this as selection on standing genetic variation where the neighboring sites can accumulate genetic diversity. This minimal threshold is set by sampling a random number between 0.05 and 0.25. After this minimum threshold is reached, the mutation shifts to being highly beneficial. Under the ancient sweep scenarios, the mutation occurs when the populations initially split and the sweep occurred until secondary contact and is allowed to segregate neutrally since then. The simulations were run without gene flow under the assumption that the sweep mutation serves as a barrier locus.

*Recent sweep (hard, soft)*

The variations in recent sweeps are identical to that of the ancient sweep with the only difference being when the mutation occurred. Under recent sweeps the mutation occurred after secondary contact and was still under selection until the sampling occurred at generation 21,952.

### Supplementary tables

Table S1. *Sampling*. All ND2 sequences were downloaded from GenBank on 18 Nov 2021. All ddRAD sampling was conducted for this study. Abbreviations: ABTC – Australian Biological Tissue Collection, South Australian Museum, Adelaide; ANSP – Academy of Natural Sciences of Philadelphia at Drexel University; ANWC – Australian National Wildlife Collection, CSIRO, Canberra; MV – Museum Victoria, Melbourne; UWBM – University of Washington Burke Museum, Seattle. *ANWC B38427 has tissues MV Z46020 (heart), MV Z46022 (muscle), and MV Z46028 (liver); ANWC B40414 (voucher) has tissues MV Z45994 (liver)/Z45997 (muscle)/Z46030 (heart); ANWC B40415 (voucher) has tissues MV Z46003 (muscle)/Z46033 )heart)/Z46038 (liver); MV ET39 is a field number matched to tissues MV Z46069 (muscle) and MV Z46031 (liver); MV ET 38 has tissues MV Z46025 (liver) and MV Z46013 (muscle); MVW031 has tissues MV Z45977 (muscle), MV Z46571 (liver); MVW038 has tissues MV Z46567 (muscle), MV Z46576 (liver). The + symbol after *A. personatus* and *A. superciliosus* specimens in the ddRAD sampling column indicates that the ANWC database has been checked for the state of reproductive organs and an assessment made that the bird was in active reproductive condition. Some subjectivity can be associated with this for male bird when interpreting the size of testes but it is less ambiguous for female birds when an oviduct is recorded as swollen or the follicles as large and golden, for example.

ND2 sampling

| GenBank Accn | Voucher | Species |
|---|---|---|
| DQ096650.1 | ANSP10590 | *A. personatus* |
| DQ096651.1 | ANSP10600 | *A. personatus* |
| DQ096652.1 | ANSP10634 | *A. personatus* |
| DQ096653.1 | ANSP10653 | *A. personatus* |
| DQ096654.1 | ANSP12312 | *A. personatus* |
| DQ096655.1 | ANSP12321 | *A. personatus* |
| DQ096656.1 | ANSP12339 | *A. personatus* |
| DQ096657.1 | ANSP12343 | *A. personatus* |
| DQ096658.1 | ANSP12348 | *A. personatus* |
| DQ096659.1 | ANSP12360 | *A. personatus* |
| DQ096660.1 | ANSP12362 | *A. personatus* |
| DQ096661.1 | ANSP12363 | *A. personatus* |
| DQ096662.1 | ANSP12364 | *A. personatus* |
| DQ096663.1 | ANSP12365 | *A. personatus* |
| DQ096664.1 | ANSP12416 | *A. personatus* |
| DQ096665.1 | ANSP12460 | *A. personatus* |
| DQ096666.1 | ANSP12467 | *A. personatus* |
| DQ096667.1 | ANSP12468 | *A. personatus* |
| DQ096668.1 | ANSP12488 | *A. personatus* |
| DQ096669.1 | ABTC2299 | *A. personatus* |
| DQ096670.1 | ABTC24318 | *A. personatus* |
| DQ096671.1 | ABTC24657 | *A. personatus* |
| DQ096672.1 | ABTC24664 | *A. personatus* |
| DQ096673.1 | ABTC27751 | *A. personatus* |
| DQ096674.1 | ABTC2838 | *A. personatus* |
| DQ096675.1 | UWBM76533 | *A. personatus* |

ddRAD sampling

| Voucher | Species |
|---|---|
| ANWC B54117 | *A. cinereus* |
| ANWC B56614 | *A. cinereus* |
| ANWC B57555 | *A. cinereus* |
| ANWC B29165 | *A. cyanopterus* |
| ANWC B45384 | *A. cyanopterus* |
| ANWC B52827 | *A. cyanopterus* |
| ANWC B47887 | *A. leucorynchus* |
| ANWC B50622 | *A. leucorynchus* |
| ANWC B57346 | *A. leucorynchus* |
| ANWC B33245 | *A. minor* |
| ANWC B55145 | *A. minor* |
| ANWC B57421 | *A. minor* |
| ANWC B28926+ | *A. personatus* |
| ANWC B28927+ | *A. personatus* |
| ANWC B28928 | *A. personatus* |
| ANWC B28986+ | *A. personatus* |
| ANWC B31976 | *A. personatus* |
| ANWC B34838 | *A. personatus* |
| ANWC B34842 | *A. personatus* |
| ANWC B48972+ | *A. personatus* |
| ANWC B49368+ | *A. personatus* |
| ANWC B49369+ | *A. personatus* |
| ANWC B49864+ | *A. personatus* |
| ANWC B49865+ | *A. personatus* |
| ANWC B55460 | *A. personatus* |
| ANWC B55461 | *A. personatus* |

| | | |
|---|---|---|
| DQ096676.1 | UWBM76535 | *A. personatus* |
| DQ096677.1 | MVET39* | *A. personatus* |
| DQ096678.1 | MVW031* | *A. personatus* |
| DQ096679.1 | MVW038* | *A. personatus* |
| DQ096680.1 | ANWC B38427* | *A. personatus* |
| DQ096681.1 | UWBM57686 | *A. personatus* |
| DQ096682.1 | UWBM57690 | *A. personatus* |
| DQ096683.1 | UWBM57691 | *A. personatus* |
| DQ096711.1 | ANWC B28926 | *A. personatus* |
| DQ096713.1 | ANWC B48972 | *A. personatus* |
| DQ096716.1 | ANWC B31976 | *A. personatus* |
| DQ096717.1 | ANWC B49369 | *A. personatus* |
| DQ096718.1 | ANWC B49368 | *A. personatus* |
| DQ096719.1 | ANWC B49864 | *A. personatus* |
| DQ096721.1 | ANWC B49865 | *A. personatus* |
| DQ096722.1 | ANWC B28928 | *A. personatus* |
| DQ096724.1 | ANWC B28927 | *A. personatus* |
| DQ096726.1 | ANWC B28986 | *A. personatus* |
| DQ096684.1 | ANSP10701 | *A. superciliosus* |
| DQ096685.1 | ANSP10704 | *A. superciliosus* |
| DQ096686.1 | ANSP10707 | *A. superciliosus* |
| DQ096687.1 | ANSP12310 | *A. superciliosus* |
| DQ096688.1 | ANSP12317 | *A. superciliosus* |
| DQ096689.1 | ANSP12318 | *A. superciliosus* |
| DQ096690.1 | ANSP12320 | *A. superciliosus* |
| DQ096691.1 | ANSP12366 | *A. superciliosus* |
| DQ096692.1 | ANSP12367 | *A. superciliosus* |
| DQ096693.1 | ANSP12370 | *A. superciliosus* |
| DQ096694.1 | ANSP12486 | *A. superciliosus* |
| DQ096695.1 | ABTC22038 | *A. superciliosus* |
| DQ096696.1 | ABTC2298 | *A. superciliosus* |
| DQ096697.1 | ABTC2410 | *A. superciliosus* |
| DQ096698.1 | ABTC24297 | *A. superciliosus* |
| DQ096699.1 | ABTC2547 | *A. superciliosus* |
| DQ096700.1 | ABTC27749 | *A. superciliosus* |
| DQ096701.1 | ABTC2985 | *A. superciliosus* |
| DQ096702.1 | UWBM76504 | *A. superciliosus* |
| DQ096703.1 | UWBM76595 | *A. superciliosus* |
| DQ096704.1 | UWBM76597 | *A. superciliosus* |
| DQ096705.1 | ANWC B40414* | *A. superciliosus* |
| DQ096706.1 | ANWC B40415* | *A. superciliosus* |
| DQ096707.1 | MVET38* | *A. superciliosus* |
| DQ096708.1 | UWBM57687 | *A. superciliosus* |
| DQ096709.1 | UWBM57688 | *A. superciliosus* |
| DQ096710.1 | UWBM57689 | *A. superciliosus* |
| DQ096712.1 | ANWC B31975 | *A. superciliosus* |
| DQ096714.1 | ANWC B44031 | *A. superciliosus* |
| DQ096715.1 | ANWC B20997 | *A. superciliosus* |
| DQ096720.1 | ANWC B32024 | *A. superciliosus* |
| DQ096723.1 | ANWC B20998 | *A. superciliosus* |

| | |
|---|---|
| ANWC B55462 | *A. personatus* |
| ANWC B55628+ | *A. personatus* |
| ANWC B55629+ | *A. personatus* |
| ANWC B55646+ | *A. personatus* |
| ANWC B55647+ | *A. personatus* |
| ANWC B55648+ | *A. personatus* |
| ANWC B57425 | *A. personatus* |
| ANWC B57426 | *A. personatus* |
| ANWC B57427 | *A. personatus* |
| ANWC B20997 | *A. superciliosus* |
| ANWC B20998 | *A. superciliosus* |
| ANWC B31975+ | *A. superciliosus* |
| ANWC B31977+ | *A. superciliosus* |
| ANWC B32024+ | *A. superciliosus* |
| ANWC B34756 | *A. superciliosus* |
| ANWC B34806 | *A. superciliosus* |
| ANWC B44031+ | *A. superciliosus* |
| ANWC B53197 | *A. superciliosus* |
| ANWC B55611 | *A. superciliosus* |
| ANWC B55612+ | *A. superciliosus* |
| ANWC B55613+ | *A. superciliosus* |
| ANWC B55614 | *A. superciliosus* |
| ANWC B57329 | *A. superciliosus* |

| | | |
|---|---|---|
| DQ096725.1 | ANWC B31977 | *A. superciliosus* |

Table S2. *Chromosome nomenclature*. Although the reads were mapped onto the *Corvus moneduloides* genome assembly, the chromosome nomenclature used in this study was based on the *Taeniopygia guttata* assembly for consistency with other studies. NOTE: The *C. moneduloides* chromosome assemblies were not reoriented to match the *T. guttata* chromosomes.

| bCorMon1.pri | TaeGut 3.2.4 |
|---|---|
| chr1 | chr2 |
| chr2 | chr1 |
| chr3 | chr3 |
| chr4 | chr1A |
| chr5 | chr4 |
| chr6 | chr5 |
| chr7 | chr7 |
| chr8 | chr6 |
| chr9 | chr8 |
| chr10 | chr9 |
| chr11 | chr12 |
| chr12 | chr11 |
| chr13 | chr10 |
| chr14 | chr4A |
| chr15 | chr13 |
| chr16 | chr14 |
| chr17 | chr20 |
| chr18 | chr15 |
| chr19 | chr18 |
| chr20 | chr19 |
| chr21 | chr17 |
| chr22 | chr21 |
| chr23 | chr23 |
| chr24 | chr26 |
| chr25 | chr24 |
| chr26 | chr27 |
| chr27 | chr22 |
| chr28 | chr28 |
| chr29 | chr25 |
| chr30 | chr29 |
| chrZ | chrZ |

Table S3. *Genes adjacent to outliers*. Genes listed here can be found 40kb upstream or downstream of Fst outliers. Alternate row shading represents a different outlier loci except for SOX5 which are four outliers clustering in one gene.

| Chrom | Start | End | CorMon Gene ID | Gene name in zebra finch | |
|---|---|---|---|---|---|
| chr1 | 28939991 | 28944204 | ENSCMUG00005005629 | GPR162 | G protein-coupled receptor 162 |
| chr1 | 28945453 | 28956176 | ENSCMUG00005005639 | P3H3 | prolyl 3-hydroxylase 3 |
| chr1 | 28963040 | 28969354 | ENSCMUG00005005657 | GNB3 | prolyl 3-hydroxylase 3 |
| chr1 | 28970031 | 28972524 | ENSCMUG00005005671 | CDCA3 | cell division cycle associated 3 |
| chr1 | 28972596 | 28990503 | ENSCMUG00005005694 | USP5 | ubiquitin specific peptidase 5 |
| chr1 | 28994046 | 28999391 | ENSCMUG00005005733 | LRRC23 | leucine rich repeat containing 23 |
| **chr1A** | **737366** | **1199125** | **ENSCMUG00005008141** | **SOX5** | **SRY-box 5** |
| chr1A | 14660946 | 14664432 | ENSCMUG00005001283 | FGF6 | fibroblast growth factor 6 |
| chr1A | 14677808 | 14681234 | ENSCMUG00005001291 | FGF23 | fibroblast growth factor 23 |
| chr1A | 14688720 | 14698580 | ENSCMUG00005001301 | TIGAR | TP53 induced glycolysis regulatory phosphatase |
| chr1A | 70715146 | 70723605 | ENSCMUG00005001086 | ATP5C1 | ATP synthase, H+ transporting, mitochondrial F1 complex, gamma polypeptide 1 |
| chr1A | 70723036 | 70734113 | ENSCMUG00005001109 | KIN | Kin17 DNA and RNA binding protein |
| chr2 | 64771152 | 64788530 | ENSCMUG00005006549 | PTER | phosphotriesterase related |
| chr2 | 64791669 | 64796976 | ENSCMUG00005006556 | C1qlr3 | complement C1q like 3 |
| chr3 | 2678414 | 2694948 | ENSCMUG00005005760 | ? | |
| chr4 | 30379051 | 30400474 | ENSCMUG00005009411 | PDGFRL | platelet derived growth factor receptor like |
| chr5 | 23968958 | 23978171 | ENSCMUG00005003009 | ALDH6A1 | aldehyde dehydrogenase 6 family member A1 |
| chr5 | 23989774 | 24011354 | ENSCMUG00005003116 | ENTPD5 | ectonucleoside triphosphate diphosphohydrolase 5 |
| chr5 | 24007689 | 24016882 | ENSCMUG00005003179 | COQ6 | coenzyme Q6, monooxygenase |
| chr5 | 24016950 | 24024967 | ENSCMUG00005003195 | FAM161B | family with sequence similarity 161, member B |
| chr13 | 6386126 | 6405398 | ENSCMUG00005012057 | G3BP1 | G3BP stress granule assembly factor 1 |
| chr13 | 6408717 | 6410221 | ENSCMUG00005012092 | ATOX1 | antioxidant 1 copper chaperone |
| chr13 | 9909467 | 9943748 | ENSCMUG00005007857 | ? | |
| **chr14** | **14899535** | **14967429** | **ENSCMUG00005006667** | **AXIN1** | **axin 1** |
| chr14 | 14968722 | 14974752 | ENSCMUG00005006676 | PDIA2 | protein disulfide isomerase family A member 2 |
| chr18 | 7810683 | 7825544 | ENSCMUG00005007292 | SRP68 | signal recognition particle 68 |
| chr18 | 7828547 | 7831702 | ENSCMUG00005007318 | GALR2 | galanin receptor 2 |
| chr18 | 7833871 | 7857101 | ENSCMUG00005007354 | EXOC7 | exocyst complex component 7 |
| chr22 | 4524264 | 4528824 | ENSCMUG00005010510 | PLPP5 | phospholipid phosphatase 5 |
| chr22 | 4534192 | 4566010 | ENSCMUG00005010547 | NSD3 | Wolf-Hirschhorn syndrome candidate 1-like 1 |
| chr22 | 4566329 | 4573492 | ENSCMUG00005010653 | LETM2 | Leucine zipper and EF-hand containing transmembrane protein 2 |
| chrZ | 24734588 | 24744717 | ENSCMUG00005013536 | CRHBP | corticotropin releasing hormone binding protein |
| chrZ | 24756488 | 24757944 | ENSCMUG00005013550 | S100Z | S100 calcium binding protein Z |
| chrZ | 53524805 | 53531087 | ENSCMUG00005015543 | NMRK1 | nicotinamide riboside kinase 1 |
| chrZ | 53533363 | 53551998 | ENSCMUG00005015550 | OSTF1 | osteoclast stimulating factor 1 |

#### Supplementary Figures

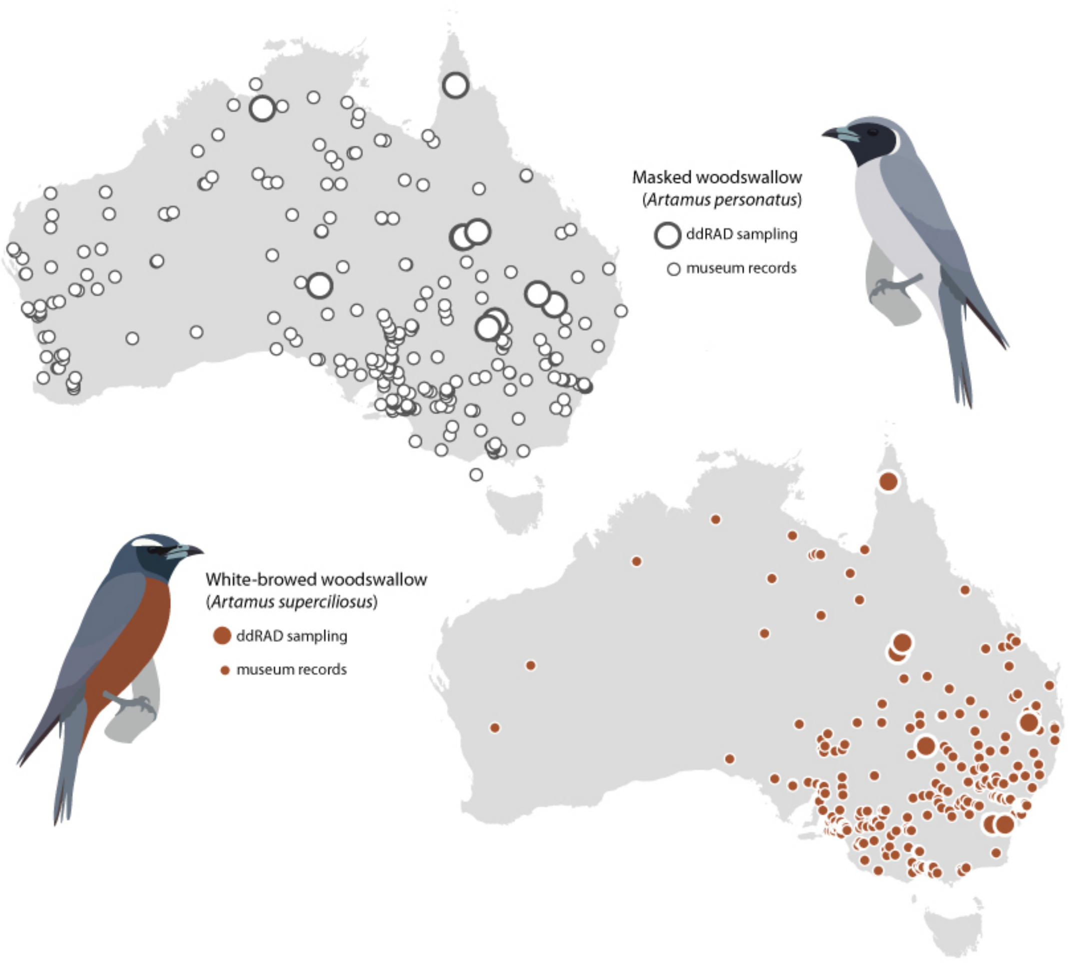


**Figure S1.** *Sampling and occurrence records.* The ddRAD sampling in this study is represented by the larger circles. The smaller circles points represent occurrence records based on museum vouchers from the Atlas of Living Australia database accessed on 24 Nov 2021.

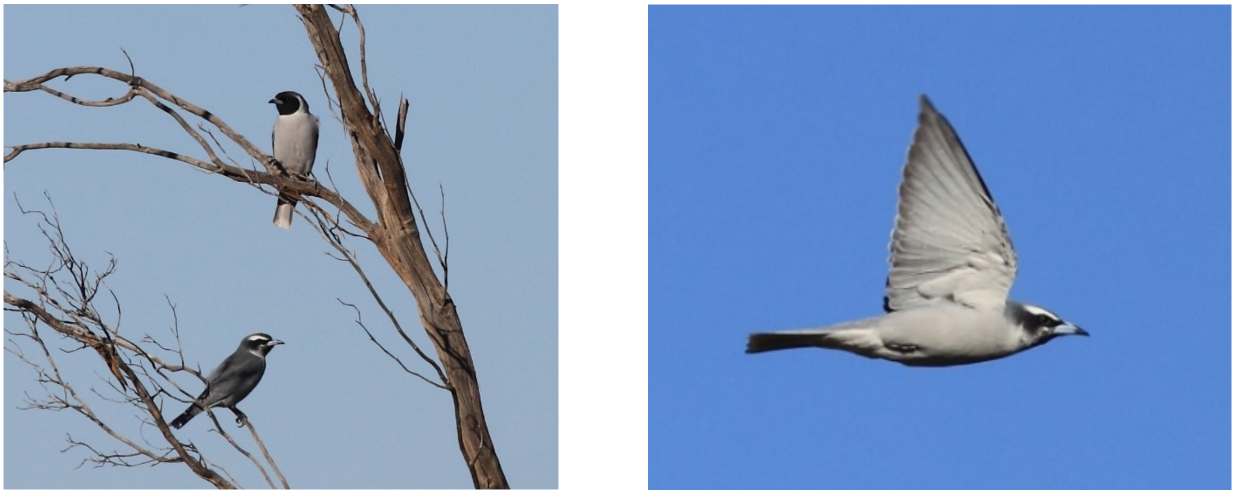

**Figure S2**. Putative hybrid (lower bird at left). Silver City Highway, 112 kilometres south of Tibooburra, New South Wales (-32.279, 142.055 (=30°16’47’’S, 142°03’17’’E)), New South Wales. 11 April 2012. Photos: Andrew Silcocks. Reproduced here with the photographer’s permission.

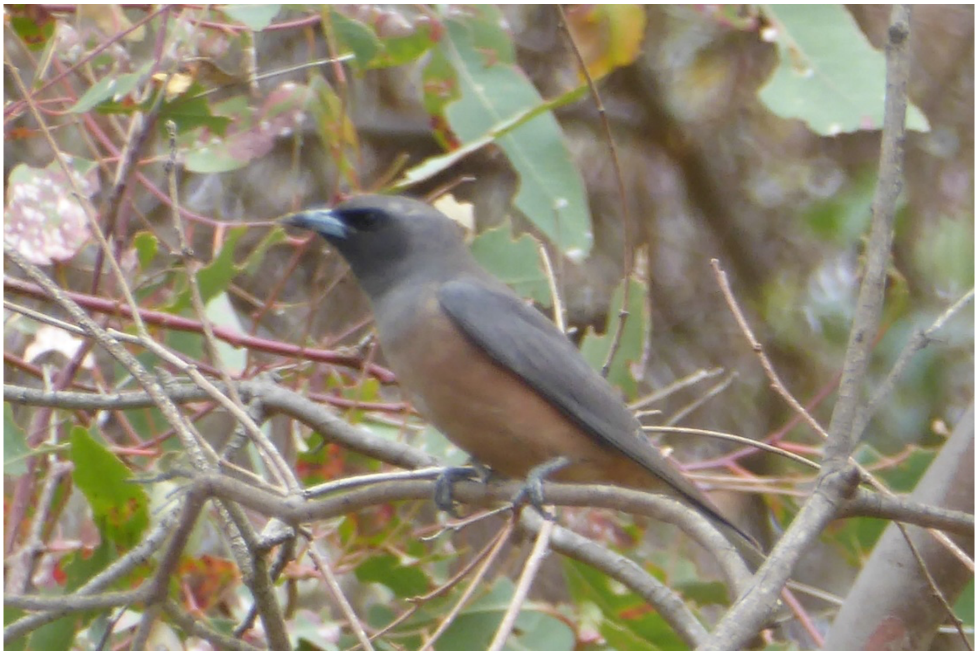

**Figure S3**. Putative hybrid. Hall Travelling Stock Reserve, New South Wales Reserve (-35.161, 149.069), 16 January 2016. Photo: Christine Darwood. Reproduced here with the photographer’s permission.

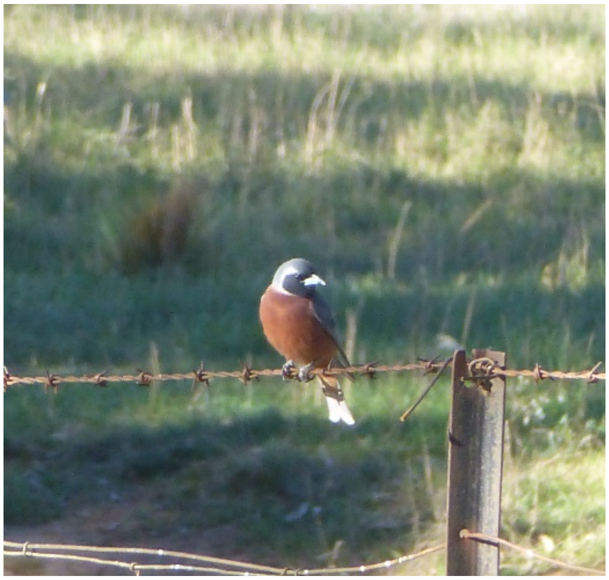

**Figure S4**. Putative hybrid. Campbell Park, Australian Capital Territory (-35.278, 149.176), 15 October 2013. Photo: Christine Darwood. Reproduced here with the photographer’s permission.

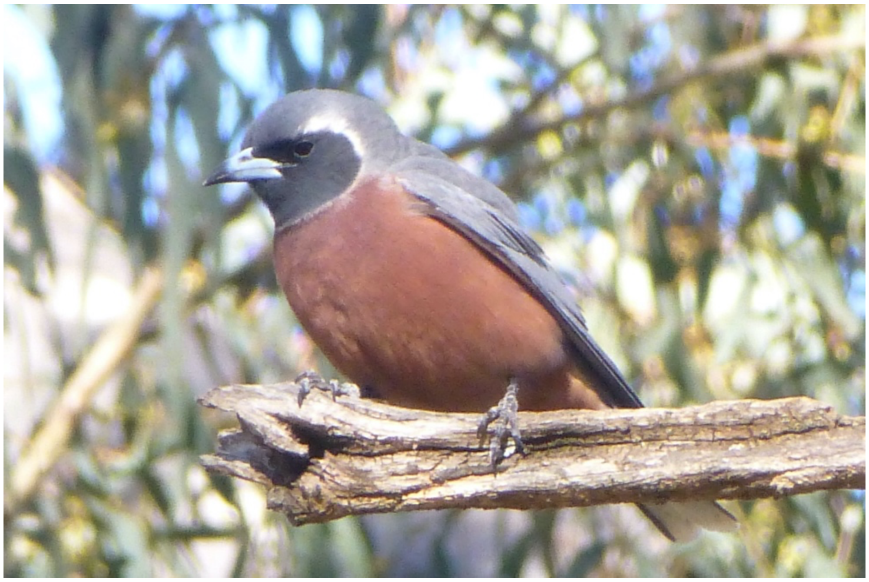

**Figure S5**. Putative hybrid. Campbell Park, Australian Capital Territory (-35.278, 149.176), 15 October 2013. Photo: Christine Darwood. Reproduced here with the photographer’s permission.

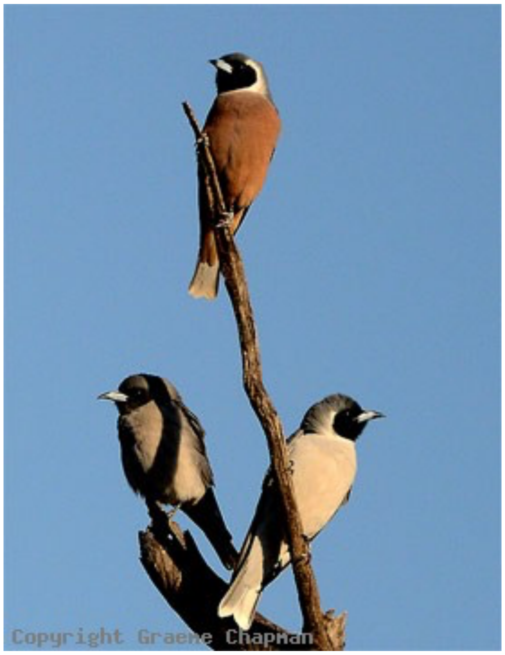


**Figure S6**. Putative hybrid. Paddington Station, southwest of Cobar, New South Wales (-32.16; 145.187), 23 September 2012.
Photo: Graeme Chapman. Reproduced here with the photographer's permission.

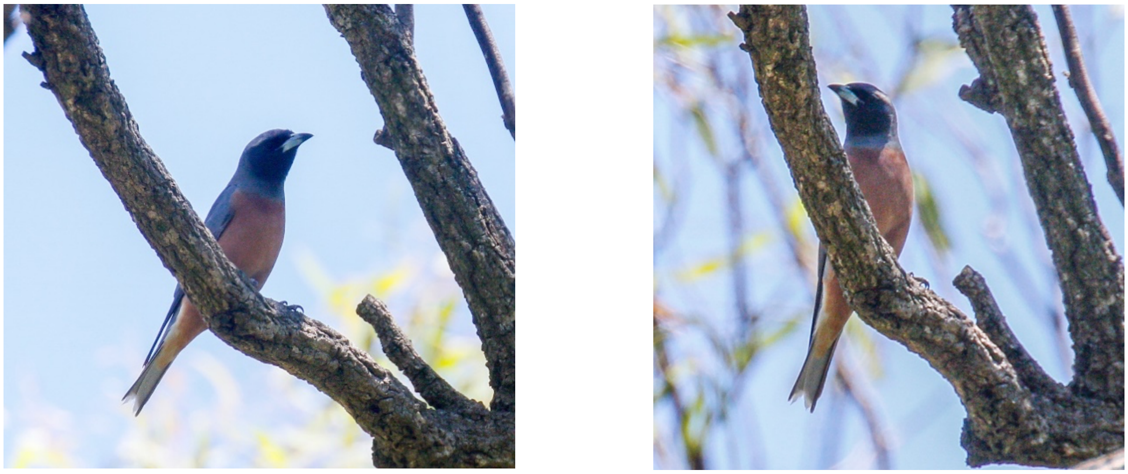

**Figure S7**. Putative hybrid. Durridgere (-32.10, 149.89), New South Wales, 14 November 2020.
Photographer: Mick Roderick. Reproduced here with the photographer's permission.

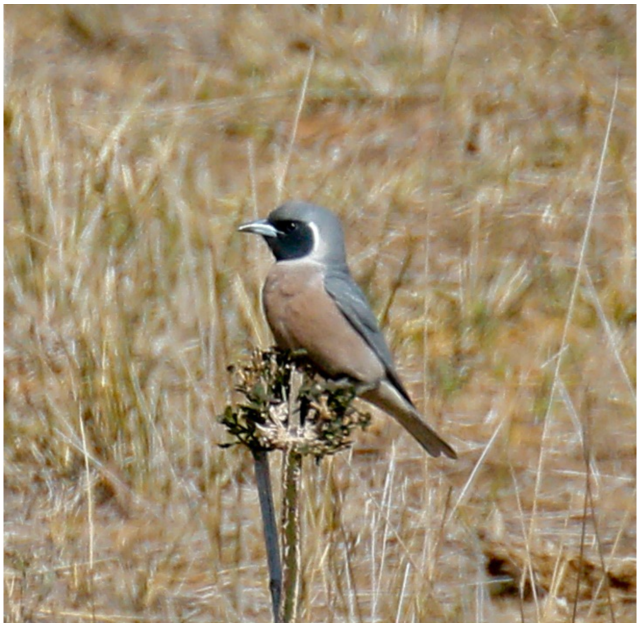

**Figure S8**. Putative hybrid. Cressfield (-31.96, 150.86), New South Wales, 28 September 2018. Photographer Mick Roderick. Reproduced here with the photographer's permission.

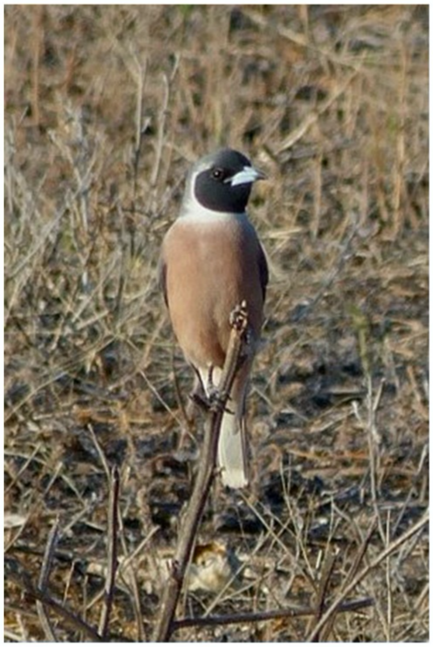

**Figure S9.** Putative hybrid. Oolambeyan National Park, ~45 kilometres south-east of Hay, New South Wales, November 2012 (further precision in location unavailable). Photo: Bastiaan Hensen. Reproduced here with the photographer's permission.

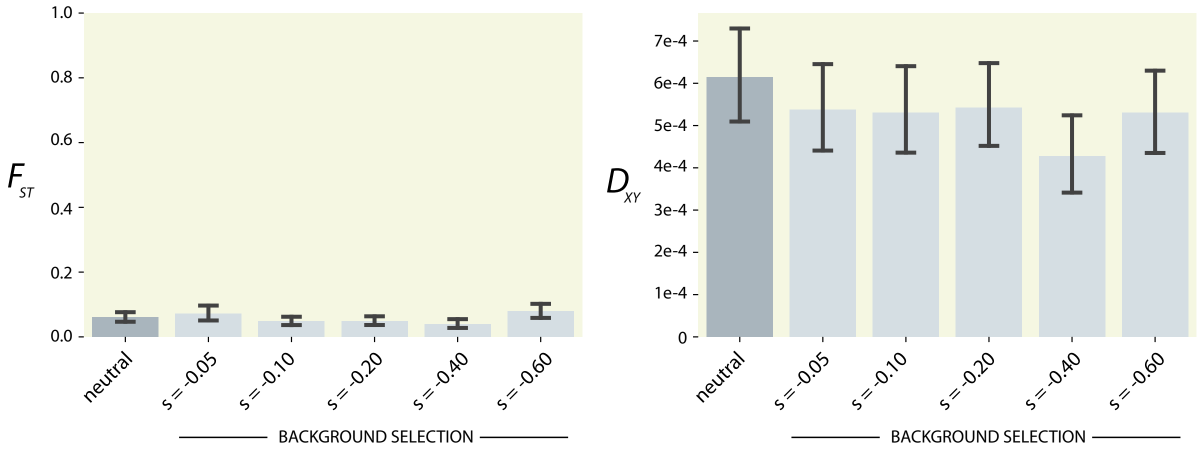


**Figure S10.** *Background selection.* The effect of different levels of background selection on relative and absolute divergence from the SLiM simulations.
